# Realizing ‘integral control’ in living cells: How to overcome leaky integration due to dilution?

**DOI:** 10.1101/141051

**Authors:** Yili Qian, Domitilla Del Vecchio

## Abstract

A major problem in the design of synthetic genetic circuits is robustness to perturbations and uncertainty. Because of this, there have been significant efforts in recent years in finding approaches to implement integral control in genetic circuits. Integral controllers have the unique ability to make the output of a process adapt perfectly to disturbances. However, implementing an integral controller is challenging in living cells. This is because a key aspect of any integral controller is a “memory” element that stores the accumulation (integral) of the error between the output and its desired set-point. The ability to realize such a memory element in living cells is fundamentally challenged by the fact that all biomolecules dilute as cells grow, resulting in a “leaky” memory that gradually fades away. As a consequence, the adaptation property is lost. Here, we propose a general principle for designing integral controllers such that the performance is practically unaffected by dilution. In particular, we mathematically prove that if the reactions implementing the integral controller are all much faster than dilution, then the adaptation error due to integration leakiness becomes negligible. We exemplify this design principle with two synthetic genetic circuits aimed at reaching adaptation of gene expression to fluctuations in cellular resources. Our results provide concrete guidance on the biomolecular processes that are most appropriate for implementing integral controllers in living cells.

## 1 Introduction

A long-standing challenge in synthetic biology is the creation of genetic circuits that perform predictably and reliably in living cells [1, 2]. Variations in the environment [3], unforeseen interactions between a circuit and the host machinery [4], and unknown interference between circuit components [5] can all significantly alter the intended behavior of a genetic circuit. Integral control, which is ubiquitous in traditional engineering systems, is a promising approach to mitigate the impact of such unknowns on genetic circuits. At the most basic level, an integral controller ensures that the output of a system can reach a desired set-point while perfectly rejecting (i.e., adapting to) constant disturbances [6–8]. Due to this unique ability, synthetic biology research has witnessed increasing efforts toward establishing procedures to realize integral control in living cells [9–17].

At the core of an integral controller is an indispensable “memory” element that accumulates (i.e., integrates) the error between the desired set-point and the measured output over time. While computing this integral is almost trivial using electronic components, realizing it through biomolecular reactions in living cells is problematic. This is because information about the error is usually stored in the form of molecular concentrations, and as host cell grows and divides, biomolecules in every single cell dilute [18, 19], leading to a “leaky” memory that fades away in time. Since cell growth is unavoidable and even beneficial in a number of applications, such as in biofuel production [20], leaky integration is a fundamental physical limitation for the implementation of integral control through genetic circuits in living cells.

Prior theoretical studies have proposed integral control motifs (ICMs), but neglected the presence of molecule dilution in the analysis [10, 12, 15, 17]. Consequently, while these motifs have theoretically guaranteed performance in cell-free systems, where dilution is non-existent, their performance in the presence of molecule dilution is not preserved. In fact, the deteriorating effect of leaky integration can be significant, as shown in this paper and in other recent studies (e.g., [21, 22]). Therefore, possible solutions to leaky integration have been proposed [10, 21]. Specifically, the approach proposed in [10] hinges on “canceling” the effect of dilution by producing the “memory species” at the same rate as its dilution. This approach, however, relies on exact parameter matching and is hence difficult to realize in practice. A different approach is to use exhaustive numerical simulation to extract circuit parameters that minimize the effects of leaky integration [21], leading to a lengthy and ad hoc design process.

In this report, we propose general design principles for ICMs that perform well despite the presence of dilution. In particular, we mathematically demonstrate that the undesirable effect of dilution can be arbitrarily suppressed by increasing the rate of all controller reactions. This is true under mild conditions that are largely independent of specific parameter values. This design principle guides the choice of core biomolecular processes and circuit parameters that are most suitable for realizing an ICM in living cells. We illustrate this guided choice on two circuits that are designed to mitigate the effects of transcription/translation resource fluctuations on gene expression, a problem that has gained significant attention recently [4, 23–25].

## 2 Quasi-integral control

The approach that we take in this paper is as follows. We first describe two types of ideal ICMs in Section 2.1, which were previously proposed abstract circuit motifs for adapting to constant disturbances in the absence of dilution. We then introduce leaky ICMs, which add dilution to these ideal ICMs, and demonstrate that the adaptation property is lost. Finally, we describe quasi-ICMs in Section 2.2, the main novelty of this paper, in which the controller reactions are engineered to be much faster than dilution, enabling the circuit to restore almost perfect adaptation to constant disturbances.

### 2.1 Ideal ICMs and leaky ICMs

We illustrate in Figure 1A two different types of ideal ICMs that abstract the two main mechanisms for biomolecular integral control proposed in the literature. In both types of motifs, we denote by *x* the species whose concentration needs to be kept at a set-point *u*, despite that its production rate is affected by a constant disturbance *d*. Therefore, the output of the process to be controlled is the concentration of species x, which is denoted by *x* (*italic*). The purpose of implementing an integral controller is thus to have the steady state of x, denoted by 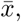 to satisfy 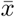 = u regardless of d (i.e., *x* adapts perfectly to d).

**Figure 1:**
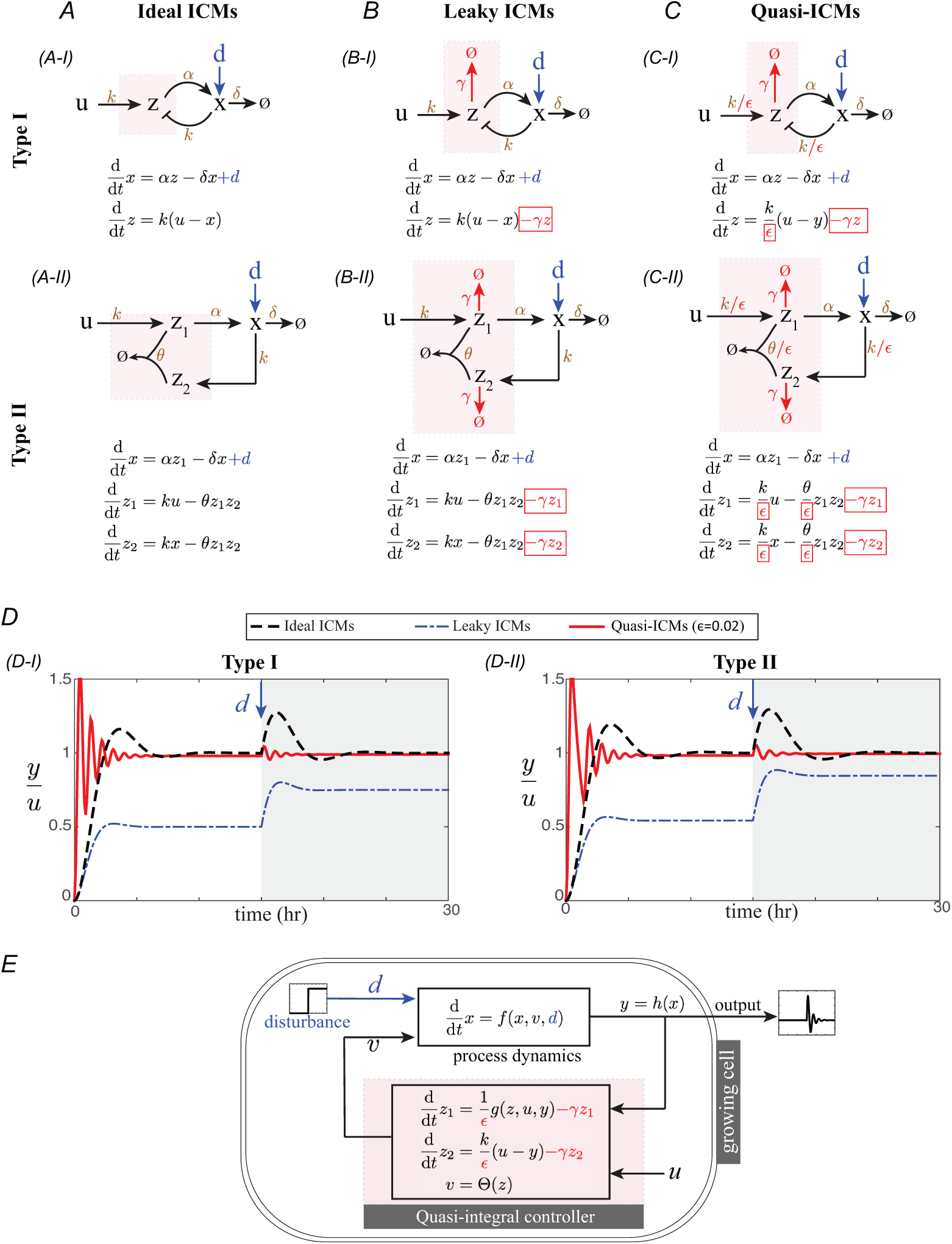
Quasi-integral control mitigates the effect of leaky integration due to dilution. (A) Two types of ideal ICMs. The controller reactions are boxed in pink, and the rest of the circuit belongs to the process to be controlled. Dilution of the controller species are neglected in ideal ICMs. (B) When dilution of the controller species are considered, ideal ICMs become leaky ICMs that cannot carry out integration. The adaptation error, defined as the extent to which the output is affected by the disturbance, can be arbitrarily large. (C) All controller reactions in a quasi-ICM are 1/∊ times faster than those in the corresponding leaky ICM, with ∊≪ 1. (D) Simulations of the type I and type II ideal, leaky and quasi-ICMs. Simulation parameters for both motifs: *α = γ = δ = k = 1* hr^−1^, *θ = 1* nM^−1^·hr^−1^, ∊ *=*0.02, *u =* 10 nM and *d =* 5 nM·hr^−1^. Disturbance input *d* is applied at 15 hr. (E) A general ∊-quasi-integral control system. Output of the process ***y*** becomes an input to the quasi-integral controller. Variable *z*_2_ is the leaky memory variable, and *z_1_* represents the remaining controller states, if any. All controller reactions are much faster than dilution, as characterized by parameter ∊. Output of the controller *v* drives the process to set-point *u* and adapt to *d*.

In these ideal ICMs, selected controller species realize the memory function essential for integral control. Specifically, a type I ideal ICM (Figure 1A-I) regulates the expression of *x* using a single controller species z. This motif arises from saturating certain Hill-type or Michaelis-Menten-type kinetics such that the production and the removal rates of z become roughly proportional to u and *x*, respectively, resulting in the following approximate mass action kinetics model for the controller species z [10, 15, 17]:

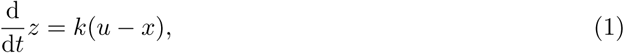

where *k* is a positive constant called integral *gain*. Since *z = ∫ k(u - x)dt, z* is called a *memory variable* that represents the integral of the error (*u - x*) between the set-point and the output over time. Independent of the exact reactants implementing this ideal ICM, it is immediate from (1) that by setting the time derivative to 0, the steady state output satisfies 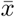 *= u* regardless of *d.*

With reference to Figure 1A-II, a type II ideal ICM, arising from what was called the antithetic integral controller in [12], realizes integral action using two controller species z1 and z2. Their production rates are engineered to be proportional to the set-point (*u*) and to the concentration of the output species (x), respectively. The two controller species can bind together to form a complex that is removed with rate constant 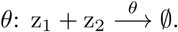 These biomolecular processes can be described by the following mass action kinetics model:

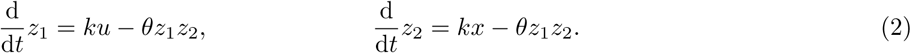

The memory function is carried out by the hidden memory variable *z* ≔ *z_1_ - z_2_*, which satisfies *dz/dt = dz_1_/dt - dz_2_/dt = k(u-x*). As a consequence, independent of the exact choice of reactants, parameters and the magnitude of disturbance *d*, at steady state, we must have 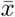 *= u*. With reference to the simulation results in Figure 1D (black dashed lines), the output *x* of both types of ideal ICMs reaches the set-point *u* and adapts to disturbance *d* perfectly at steady state.

When dilution of the controller species due to host cell growth is taken into account, the key integral structure of the memory variables is disrupted (Figure 1B). In fact, in a standard mass action kinetics model describing reactions in exponentially growing cells, the average effect of cell growth and division on the dynamics of any species z can be modeled by a first-order additive term -γz, where γ is the specific growth rate of the host cell [18, 19, 26, 27] (see SI Section S1.1 for details). Hence, with reference to Figure 1B-I, the dynamics of the memory variable *z* in a type I ICM become

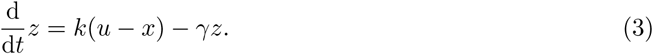

Similarly, as shown in Figure 1B-II, the dynamics of the two controller species z_1_ and z_2_ in the type II ICM become

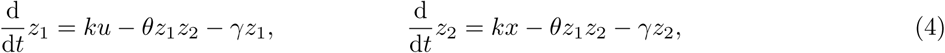

resulting in the dynamics of the hidden memory variable to become

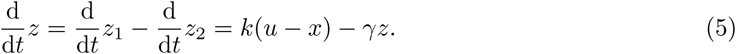

From equations (3) and (5), we observe that in both types of motifs, the memory variable *z* is no longer integrating the error between the set-point and the output, but rather carries out leaky integration. We therefore call the motifs in Figure 1B *leaky ICMs*. In simulations of Figure 1D (blue dash-dot lines), we demonstrate that including dilution significantly hinders the ability of leaky ICMs to achieve adaptation to disturbance *d*, resulting in nearly 100% relative adaptation error. More generally, as we exemplify in SI Section S1.5, for any fixed dilution rate constant (γ), depending on the process to be controlled and the parameters in the leaky ICM, the adaptation error can be arbitrarily large.

### 2.2 Quasi-ICMs: almost perfect adaptation to disturbances

In this section, we introduce quasi-ICMs, the main contribution of this paper. While leaky ICMs cannot achieve perfect adaptation to disturbances, and their performance can be arbitrarily deteriorated by the presence of dilution, we propose that one can achieve almost perfect adaptation to disturbances despite dilution by increasing the rate of all controller reactions in the leaky ICMs. Therefore, we call a leaky ICM whose controller reactions are much faster than the slower dilution process a *quasi-ICM*. These motifs are shown in Figures 1C-I and C-II respectively for the two types of motifs. In particular, with reference to Figure 1C-I, we use a small dimensionless positive parameter 0 < ∊ ≪ 1 to capture the fact that the controller reaction rates in a type I quasi-ICM are 1/e times faster than those in a type I leaky ICM (Figure 1B-I), resulting in the memory variable dynamics to become

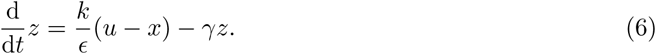

Similarly, when all the controller reaction rates in a type II leaky ICM are increased by a factor of 1/∊ (see Figure 1C-II), the controller species dynamics become

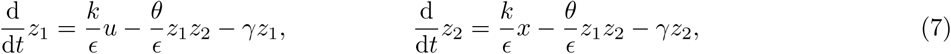

resulting in the following hidden memory variable dynamics:

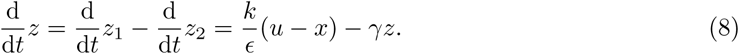

As shown in Figure 1D (red solid lines), for small ∊, both types of quasi-ICMs restore the ability to drive the output *x* to the set-point *u*, and to adapt to *d* almost perfectly despite dilution. These qualitative observations are reflected mathematically in equations (6) and (8), where for both motifs the steady state adaptation error e can be computed as e = *u* - 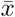 *= ∊γ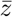/k*, whose magnitude can be arbitrarily decreased by detuning ∊ (i.e., increasing controller reaction rates). Although this reasoning is intuitive, it is based on the implicit assumption that the steady state concentration of controller species *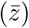* stays roughly unchanged as ∊ decreases, which is, unfortunately, not always true and hard to verify in general (see SI Section S1.6 for such an example). Therefore, in the rest of this section, we provide precise and general mathematical conditions under which performance deterioration of a leaky integral controller can be arbitrarily suppressed by faster controller reactions. These results establish a general circuit design principle applicable to any biomolecular process to be controlled and to any leaky integral controller, including, but not limited to those arising from the type I and type II ICMs.

We consider a biomolecular process to be controlled, whose dynamics can be written as:

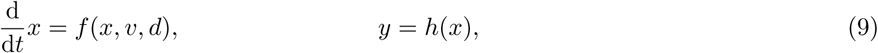

where *x* represents process states (i.e., the concentrations of species *x* forming the biomoelcular process). The process takes two inputs: *v* is the control input produced by the controller, and *d* is a disturbance input, which could also represent an uncertainty in process parameters. The output of the process *y* is determined by a function *h(x*). The process (9) is connected to an ∊-parameterized biomolecular controller, which contains a leaky integral controller. We study the case where all controller reactions have an ∊-parameterized time-scale separation from dilution. To this end, we write the controller in the general form:

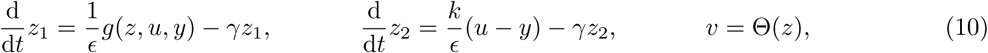

where *z* ≔ *[z_1_,z_2_]^T^* represents the controller states (e.g., concentrations of controller species) and ∊ is a small positive parameter. Specifically, *z_2_* is the intended memory variable that carries out the leaky integration with gain *k*, and *z_1_* represents concentrations of additional controller species, if any. The controller takes the output of the process *y* as input, and compares it with a set-point *u*. The *closed loop system* is therefore a feedback interconnection of process (9) and controller (10) (see Figure 1E). Assuming that the closed loop system has a *unique locally asymptotically stable steady state* 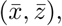 we formally define ∊-quasi-integral control as follows.

#### Definition 1.

*The closed loop system (9)-(10) realizes ∊-quasi-integral control in an admissible input set* 𝕌 x ⅅ *if for all (u*, d) ∊ 𝕌 x ⅅ, *the system’s steady state output 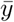 satisfies*

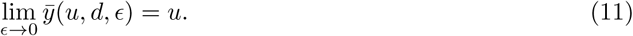

Equation (11) implies that, by increasing the rate of controller reactions (i.e., decreasing ∊), the steady state output *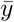* of an ∊-quasi-integral control system can be made arbitrarily close to the set-point *u* in the presence of disturbance *d*. The following claim provides an easy-to-check condition for ∊-quasi-integral control.

**Claim 1.** *The closed loop system (9)-(10) realizes ∊-quasi-integral control if the auxiliary ideal integral control system consisting of the process (9) and the ideal integral controller*

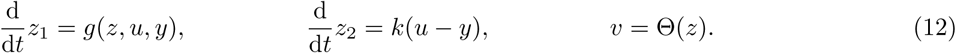

*has a unique locally exponentially stable steady state for all* (u, *d*) ∊ 𝕌 x ⅅ.

The proof of this Claim can be found in SI Section S1.2. It establishes that if one can use the ideal integral controller (12) to obtain perfect adaptation in the absence of dilution, then when dilution is taken into account, one can always speed up all controller reactions to restore adaptation. From an engineering perspective, one should therefore select controller biomolecular processes whose rates are significantly faster with respect to dilution rates. Such processes include, for example, enzymatic reactions (i.e., phosphorylation, methylation, and ubiquitination), protein-protein interactions, and RNA-based interactions (i.e., sRNA or microRNA-enabled degradation or sequestration). With these processes, the designer can then realize type I or type II ideal ICMs as well as additional ideal ICMs that are in the form of system (12). It is worth to note that even in cases where only part of the controller reactions can be sped up, it may still be possible to reach ∊-quasi-integral control, although the algebraic conditions to check become slightly more involved (see SI Section S1.7). In SI Section S4.3, we further demonstrate that when dilution rate constant varies slowly in time, the adaptation error is still guaranteed to decrease as ∊ is decreased.

Claim 1 provides the theoretical underpinning to the type I and type II quasi-ICM designs of Figure 1C. Since the corresponding type I and II ideal ICMs are in the form of (12) and have unique locally exponentially stable steady states (see proof in SI Section S1.3-1.4), increasing all controller reaction rates will quench the steady state effect of leaky integration. Furthermore, we also show in SI Section S1.3-1.4 that the steady states of the type I and II quasi-ICMs are unique and stable.

## 3 Two biomolecular realizations of quasi-ICMs

In this section, we propose physical implementations of a phosphorylation-based quasi-integral controller and a small RNA-based quasi-integral controller. They are realizations of the type I and type II quasi-ICMs, respectively, which satisfy the general design principle of fast controller reactions. For illustration purposes, we use these controllers to mitigate the effect of transcriptional and translational resource fluctuations on gene expression, a problem that has received considerable attention in synthetic biology in recent years [4, 23–25].

### 3.1 Type I quasi-ICM: phosphorylation cycle

A diagram of the phosphorylation-based quasi-integral control system is shown in Figure 2A. The controller is intended to regulate the production of protein p to adapt to a disturbance *d*, which models a reduction in protein production rate due to, for example, depletion of transcriptional and/or translational resources in the host cell [4, 23]. In this system, the intended integral control is accomplished by a phosphorylation cycle. The regulated protein p is co-expressed with a phosphatase and the concentration of the kinase acts as the set-point *u*. The substrate b is expressed constitutively. When b is phosphorylated by kinase u to become active substrate b*, it transcriptionally activates production of protein p. A simplified mathematical model of this system is (see SI Section S2.1 for detailed derivation from chemical reactions):

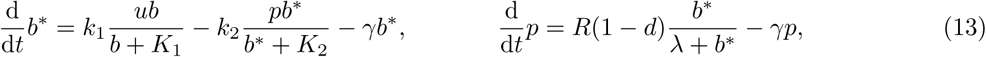

where *k_1_* and *k_2_* are the catalytic rate constants of the phosphorylation and dephosphorylation reactions, respectively, *K_1_* and *K_2_* are the Michaelis-Menten constants, *R* is the protein production rate constant, and λ is the dissociation constant between b* and the promoter of the regulated gene.

**Figure 2:**
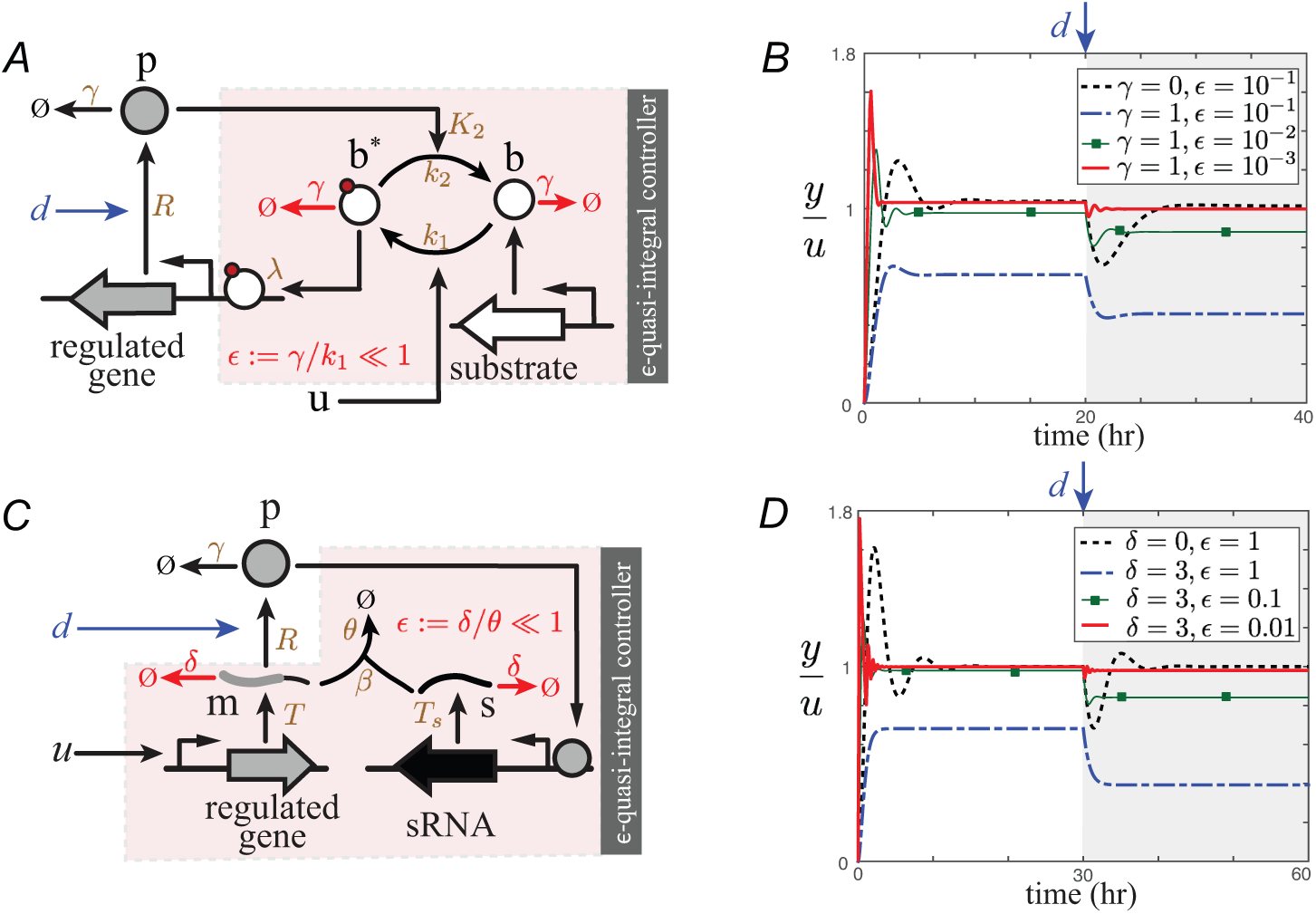
Two physical realizations of quasi-ICMs. (A) Genetic circuit diagram of the phosphorylation-based quasi-integral controller. Chemical reactions realizing the controller are boxed in pink. (B) Simulation of the circuit’s response according to (13). A set-point input *u* = 20 nM is applied at time 0 and a disturbance input *d* = 0.5 is applied at 20 hr. The vertical axis represents the ratio between output *y*, defined to be proportional to *p* (*y* = *σp*), and set-point *u*. The dashed black line is the response of the phosphorylation-based control system assuming no dilution of the active substrate b*. The dotted blue line, the thin green line with square markers and the solid red line represent circuit’s response in the presence of nonzero substrate dilution (*γ* = 1 hr^−1^) and decreasing ∊, which corresponds to increasing catalytic rates (*k*_*i*_, *i*= 1,2). (C) Genetic circuit diagram of the sRNA-based quasi-integral controller. (D) Simulation of the circuit’s response according to (16). A set-point input *u* = 1 is applied at time 0 and a disturbance input *d* = 0.5 is applied at 30 hr. The vertical axis represents the ratio between output *y*, defined in (18), and set-point input *u.* The dashed black line represents response of an ideal integral control system, where RNA decay rate *δ* is set to 0. The dotted blue line, the thin green line with square markers and the solid red line represent circuit’s responses in the presence of nonzero RNA decay rate (*δ* = 3 hr^−1^, corresponding to half-life of about 13 mins) and decreasing ∊. Parameter ∊ is decreased by increasing the mRNA-sRNA removal rate (*θ/β*). The DNA copy numbers of the regulated gene and the sRNA are increased simultaneously by a factor of 1/∊ as ∊ decreases. Simulation parameters are listed in SI Section S5, Table 2.

System (13) can be taken to the form of a type I leaky-ICM (3) if both the kinase and the phosphatase are saturated (i.e., *b* ≫ *K_1_* and *b^*^* ≫ *K_2_*). These design constraints are present in any type I ICM, which relies on saturation of Hill-type or Michaelis-Menten kinetics [10, 15, 17], and can be satisfied in practice when the substrate is over-expressed and the kinase concentration is not too small (see SI Section S2.3 for details). Under these assumptions and by setting *x* ≔ *p, z* ≔ *b^*^* and *σ* ≔ *k*_2_/*k*_1_, we can approximate (13) by

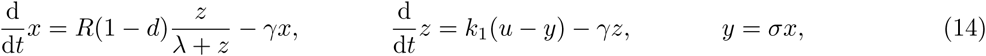

which is in the form of a type I leaky-ICM. Since phosphorylation and dephosphorayltion reactions are typically much faster than dilution, by choosing a phosphorylation cycle to realize the type I motif, the effect of leaky integration is naturally mitigated. In fact, by setting ∊≔ γ/*k*_1_ (∊ ~ 10-^3^ in bacteria [19]), system (14) is in the form of a type I quasi-ICM:

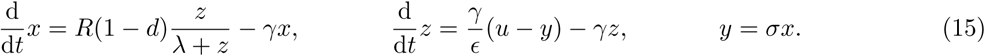

As demonstrated in the simulation results in Figure 2B and Figure S8 in SI, the steady state adaptation error due to leaky integration becomes smaller as ∊ decreases (i.e., larger time-scale separation between phosphorylation/dephosphorylation and dilution). We refer the readers to SI Section S2.2 for mathematical details on the uniqueness and stability of the steady state.

### 3.2 Type II quasi-ICM: small RNA (sRNA) interference

The sRNA-based quasi-integral controller is a realization of the type II quasi-ICM. It is intended to regulate translation of protein p to adapt to a disturbance *d*, which models uncertainty in translation rate constant due to fluctuation in the amount of available ribosomes [4, 23, 24]. With reference to Figure 2C, the controller consists of protein p transcriptionally activating production of an sRNA (s) that is complementary to the mRNA (m) of the regulated protein p. The sRNA and mRNA can bind and degrade together rapidly [28–30]. The mRNA concentration m is the control input to the translation process. A constant upstream transcription factor regulates mRNA production as a set-point input *u* to the controller. Based on the chemical reactions in SI Section S3.1, the ODE model of this system is:

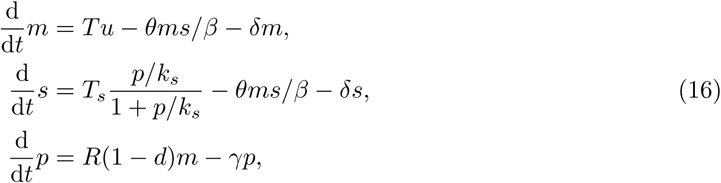

where *T* and *T_s_* characterize the production rates of mRNA and sRNA, respectively. They are proportional to the copy numbers of the regulated gene and the sRNA-expression DNA. Parameter *k_s_* is the dissociation constant of the binding between protein p and the sRNA promoter, *β* is the dissociation constant of mRNA-sRNA binding, *θ* is the degradation rate constant of the mRNA-sRNA complexes, and *R*is the translation rate constant. In addition to dilution due to cell growth, characterized by rate constant *γ*, uncoupled mRNA and sRNA are degraded by RNAse [31, 32]. Therefore, we model decay (i.e., dilution and degradation) of uncoupled RNAs by a lumped rate constant *δ* such that *δ ≥ γ*, and assume that this rate constant is the same for mRNA and sRNA without loss of generality. Our model (16) is similar to established sRNA-based translation regulation models [29, 33, 34], except that expression of sRNA is activated by the regulated protein (p) to constitute a feedback loop. These additional reactions are modeled using a standard Hill-type function in (16), and we refer the readers to SI Section S3.1 for its derivation.

Setting *z_1_* ≔ m, *z_2_* ≔ *s* and *x ≔ p*, system (16) can be taken to the form of a type II leaky-ICM:

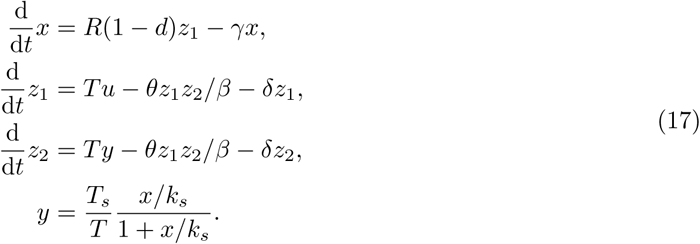

In order to achieve negligible adaptation error, we need to ensure that the controller reactions (i.e., *z*_1_ and *z*_2_ dynamics) are much faster than uncoupled RNA decay. On the one hand, coupled degradation of mRNA-sRNA complexes are by nature much more rapid than uncoupled RNA decay [30]. We therefore use *∊* ≔ *δ*/*θ* ≪1 to characterize this time-scale separation. Additionally, we can simultaneously increase DNA copy numbers of the controller species m and s by a factor of 1/e to increase their production rates. Consequently, the production rate constants of mRNA and sRNA become T/∊ and T_s_/∊, respectively. Under these assumptions, system (17) is in the form of a type II quasi-ICM:

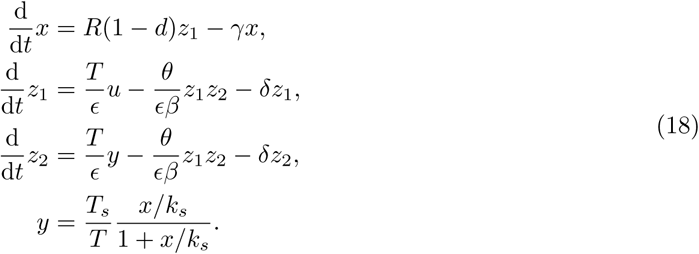

The adaptation error can be arbitrarily decreased by detuning *∊*. In practice, this can be accomplished by simultaneously (i) increasing the mRNA-sRNA complex degradation rate constant *θ*, and (ii) increasing the copy numbers of the regulated gene and the sRNA-expression DNA. While directly increasing *θ*may be difficult to implement in practice, since the parameters *θ* and *β* are clustered together in model (16), we can achieve the same effect by increasing the affinity between sRNA and mRNA (1/*β*) [35]. The above results are confirmed by simulations in Figure 2D and Figure S9 in SI using biologically relevant parameters from bacteria *E. coli.*In SI Section S3.2, we analytically demonstrate that the conditions in Claim 1 are indeed satisfied for system (18).

## 4 Discussion

Due to integral controllers’ unique ability to drive a process to reach a set-point regardless of disturbances/uncertainties, they are especially appealing to synthetic biology, where various forms of disturbances and uncertainties limit the robustness and predictability of synthetic circuits. A unique and fundamental challenge to realize integral control in living cells is that the concentration of biomolecules dilute as cells grow, resulting in the impossibility to create a non-leaky biomolecular memory element, which is indispensable to implement integral control.

In this report, we propose a general design principle to overcome this obstacle. We mathematically demonstrate that one can design an ideal integral controller to obtain perfect adaptation in the absence of dilution and then speed up all controller reactions with respect to dilution to restore adaptation in the presence of dilution. This implies that the designer should select controller biomolecular processes whose rates are significantly faster than dilution rates. Our result is equally applicable to situations where controller species are degraded in addition to being diluted. Due to the asymptotic nature of our results, they are largely insensitive to system parameters. To solidify this point, we simulated the phosphorylation-based and the sRNA-based quasi-integral controllers with randomly generated parameters, and demonstrated that, regardless of the rest of the parameters, the adaptation error always decreases as the rates of controller reactions are increased (SI Section S5.2).

While our mathematical analyses are carried out using ordinary differential equation models derived from mass action kinetics, we numerically exemplify their effectiveness using a time-varying Gillespie simulation [36], in which the stochasticity of biomolecular reactions as well as cell volume expansion and division are explicitly taken into account (see SI Section S5.3 for details). In addition, we have assumed that the host cells grow at constant rate (constant γ). In practice, however, their growth rate may change with, for example, temperature, nutrients and metabolic load of the circuit [37]. We have therefore studied the performance of quasi-integral controllers in changing growth conditions (see SI Section S4.3), and found that the circuit’s output can still adapt to disturbances as long as growth rate changes slowly.

Nature may have already utilized the time-scale separation strategy between process and controller that we propose here for realizing adaptation and homeostasis. For example, it has been suggested that the bacterial chemotaxis system [38] and the yeast osmoregulation system [39] contain integral controllers realized through methylation and phosphorylation, respectively. Since these processes are much faster than dilution in their respective host cells, it is justifiable to neglect dilution in these systems, and almost perfect adaptation can be achieved. Therefore, experimental study of synthetic realizations of the quasi-integral controllers proposed here may enhance our understanding of their natural counterparts, which are often embedded in more complex biomolecular networks and are therefore more difficult to analyze.

From a classical control-theoretic perspective, integral controllers often operate with low integral gains (i.e., slow integration). This is because with a perfect integrator, steady state adaptation can be achieved regardless of the magnitude of the integral gain, and a high integral gain often leads to undesirable effects such as higher energy consumption and tendency to instability [6, 22]. However, a potential advantage of increasing the leaky integral gain is that it can improve a system’s ability to track (reject) time-varying references (disturbances). While we demonstrate this capability for the two example systems through linearization and simulations in SI Section S4.1-4.2, further theoretical assessment of this property requires deeper study [40, 41].

## Data accessibility

All simulation codes are provided as Electronic Supplementary Material.

## Author contributions

Y.Q derived the mathematical model and performed analyses and simulations; Y.Q and D.D.V wrote the manuscript.

## Acknowledgement

We thank Cameron McBride for proof-reading the manuscript; Prof. Mustafa Khammash, Dr. Corentin Briat, Dr. Hsin-Ho Huang, Ross Jones and Theodore W. Grunberg for helpful discussions and suggestions; and Prof. Elisa Franco and Dr. Christian Cuba Samaniego for comments on earlier versions of the manuscript. We thank the anonymous reviewers for their constructive comments.

